# Mutations in an unrecognized internal NPT2A PDZ motif disrupt phosphate transport causing congenital hypophosphatemia

**DOI:** 10.1101/2023.03.06.531332

**Authors:** W Bruce Sneddon, Peter A Friedman, Tatyana Mamonova

## Abstract

The Na^+^-dependent phosphate cotransporter-2A (NPT2A, SLC34A1) is a primary regulator of extracellular phosphate homeostasis. Its most prominent structural element is a carboxy-terminal PDZ ligand that binds Na^+^/H^+^ Exchanger Regulatory Factor-1 (NHERF1, SLC9A3R1). NHERF1, a multidomain PDZ protein,establishes NPT2A membrane localization and is required for hormone-sensitive phosphate transport. NPT2A also possesses an uncharacterized internal PDZ ligand. Two recent clinical reports describe congenital hypophosphatemia in children harboring Arg^495^His or Arg^495^Cys variants within the internal PDZ motif. The wild-type internal ^494^TRL^496^ PDZ ligand binds NHERF1 PDZ2, which we consider a regulatory domain. Ablating the internal PDZ ligand with a ^494^AAA^496^ substitution blocked hormone-sensitive phosphate transport. Complementary approaches, including CRISPR/Cas9 technology, site-directed mutagenesis, confocal microscopy, and modeling, showed that NPT2A Arg^495^His or Arg^495^Cys variants do not support PTH or FGF23 action on phosphate transport. Coimmunoprecipitation experiments indicate that both variants bind NHERF1 similarly to WT NPT2A. However, in contrast to WT NPT2A, NPT2A Arg^495^His or Arg^495^Cys variants remain at the apical membrane and are not internalized in response to PTH. We predict that Cys or His substitution of the charged Arg^495^ changes the electrostatics, preventing phosphorylation of the upstream Thr^494^, interfering with phosphate uptake in response to hormone action, and inhibiting NPT2A trafficking. We advance a model wherein the carboxyterminal PDZ ligand defines apical localization NPT2A, while the internal PDZ ligand is essential for hormone-triggered phosphate transport.

## Introduction

NPT2A (SLC34A1), Na^+^-dependent phosphate cotransporter-2A, is the primary regulator of renal phosphate absorption and serum phosphate homeostasis. Disordered phosphate metabolism associated with chronic kidney disease and frank resistance to hormone action contributes to exceptionally high mortality rates, especially among the elderly and impoverished [1-3].

The most prominent NPT2A structural element is its carboxy-terminal TRL^639^ (Thr-Arg-Leu^639^) PDZ ligand, a Class I PDZ binding site (-Thr/Ser^*−*2^-X^*−*1^-Φ^0^, where X at position -1 is permissive and Φ at position 0 is any hydrophobic residue) (**Figure 1**). Through its carboxy-terminal TRL^639^ sequence, NPT2A binds Na^+^/H^+^ Exchanger Regulatory Factor-1 (NHERF1, SLC9A3R1) (Figure 1), a multidomain PDZ protein, that determines NPT2A apical localization [4-8].

**Figure 1.**
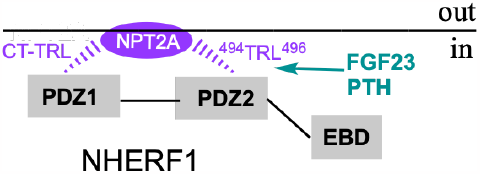
Schematic representation of NPT2A bound NHERF1 at the apical membrane. The NPT2A carboxy-terminal (CT-TRL) and internal 494TRL496 PDZ ligands bind NHERF1 PDZ1 and PDZ2, respectively. PTH-triggered phosphorylation of Thr494 in the internal 494TRL496 PDZ ligand of NPT2A is necessary for hormone-sensitive phosphate transport.

NHERF1 assembles multiprotein complexes through its two PDZ domains and an ezrin-binding domain (EBD) (Figure 1). Ser/Thr^*−*2^ and Φ^0^ contribute primarily to binding affinity. The carboxy-terminal NPT2A PDZ ligand binds NHERF1 PDZ1 [9, 10]. NHERF1 scaffolds relevant kinases [8, 11] and phosphatases [12] and controls the association and disassembly of the NHERF1:NPT2A complex, endocytosis, and cessation of phosphate transport. Mice lacking NHERF1 [13-15] and humans harboring NHERF1 mutations [16, 17] exhibit frank phosphaturia even in the presence of normal-to-high Parathyroid hormone (PTH) and Fibroblast Growth Factor 23 (FGF23) levels. PTH and FGF23, despite different membrane receptors and distinct signaling pathways, exert their shared phosphaturic action by dissociating NPT2A from NHERF1, sequestering NPT2A, thereby reducing the abundance of NPT2A at apical cell membranes and augmenting phosphate excretion [18-20].

Remarkably, in addition to its well-characterized canonical carboxy-terminal PDZ ligand, NPT2A possesses an uncharacterized internal ^494^Thr-Arg^495^-Leu^496^ PDZ sequence (Figure 1) in the intracellular loop between transmembrane domains 6 and 7. Two recent clinical reports describe inherited autosomal dominant hypophosphatemia in children harboring Arg^495^His or Arg^495^Cys variants within this internal PDZ motif [21, 22]. The presence of these mutations within a PDZ ligand was unappreciated. The role of this cryptic internal PDZ binding motif in phosphate regulation is unknown.

This study uses complementary approaches, including CRISPR/Cas9 technology, to generate an informative, hormone-sensitive cell line, site-directed mutagenesis, confocal microscopy, and modeling to characterize the wild-type and disease-associated NPT2A internal ^494^TRL^496^ PDZ motif and evaluate its participation in FGF23 and PTH-triggered phosphate regulation. We show that NPT2A Arg^495^Cys or Arg^495^His variants bind NHERF1 but do not support FGF23 or PTH action on phosphate transport. In contrast to wild-type NPT2A, Arg^495^Cys or Arg^495^His NPT2A variants remain at the apical membrane and are not endocytosed in response to PTH. We predict that Cys or His substitution of the charged Arg^495^ changes the electrostatics, preventing phosphorylation of the upstream Thr^494^, thus interfering with phosphate uptake and inhibiting NPT2A trafficking in response to hormone action. Consistent with this scheme, ablating the internal PDZ ligand blocked hormone-sensitive phosphate transport.

We advance a conceptual model where the NPT2A carboxy-terminal PDZ ligand binds NHERF1 PDZ1 and serves as a scaffold for tethering required kinases and phosphatases with apically localized NPT2A. We propose that the internal NPT2A PDZ motif is directly involved and necessary for hormone-sensitive phosphate transport via association with NHERF1 PDZ2 (Figure 1).

## Results

### The internal NPT2A ^494^TRL^496^ PDZ motif determines hormone-sensitive phosphate transport

We first interrogated whether the internal NPT2A ^494^TRL^496^ PDZ motif interacts with NHERF1. For these experiments, we developed CRISPR/Cas9 Npt2a knockout opossum kidney (OK) cells, where the native Npt2a transporter was eradicated. Deleting endogenous Npt2a allowed the introduction of wild-type human NPT2A or other designed structurally informative constructs. CRISPR/Cas9 Npt2a knockout OK cells were transfected with HA-GFP-NPT2A, where the carboxy-terminal -TRL^639^ (hereafter CT-TRL) PDZ ligand was replaced with a triple Ala sequence (hereafter CT-AAA). This modification prevents binding between NPT2A CT-TRL and NHERF1 PDZ1 domain [9, 10]. We used WT FLAG-NHERF1 or NHERF1 constructs harboring carboxylate-binding loop GYGF to GAGA modifications in PDZ1 (S1), PDZ2 (S2), or both PDZ domains (S1S2) that disrupt the formation of the canonical interactions between PDZ domains and carboxy-termini of the target ligands [9, 11, 23]. Coimmunoprecipitation data show that the NPT2A CT-AAA mutant associates with NHERF1 S1 (PDZ2) but not with S2 (PDZ1) or S1S2, indicating that the NPT2A internal ^494^TRL^496^ PDZ motif binds NHERF1 PDZ2 (**Figure 2**).

**Figure 2.**
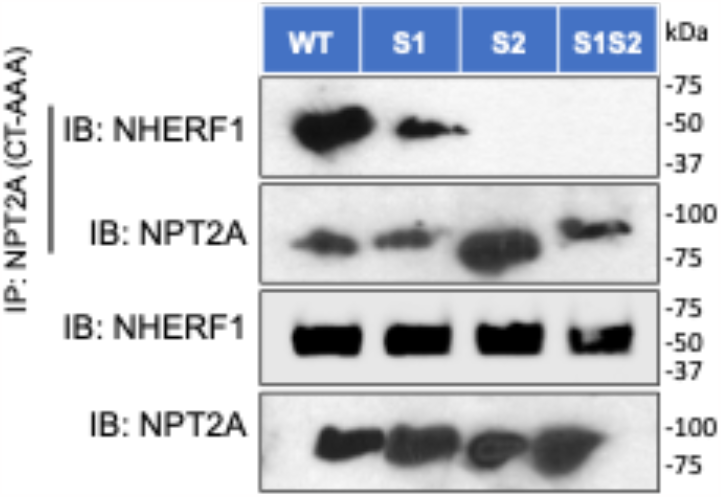
Immunoprecipitation of NPT2A with NHERF1. NPT2A with triple Ala replacement at the Molecular weight carboxy-terminal PDZ ligand(CT-AAA) immunoprecipitated (IP) with NHERF1 S1 (PDZ2) but not with S2 (PDZ1)or S1S2, indicating the NPT2A internal ^494^TRL^496^ PDZ ligand binds NHERF1 PDZ2. CRISPR/Cas9 Npt2a knockout OK cells were transfected with HA-GFP-NPT2A CT-AAA and FLAG-NHERF1 (WT, S1, S2, S1S2). Immunoblots for IP are depicted in the top two panels. Lysates controls are in the bottom panels. Molecular weight markers (kDa) on the right of blots. *n*=3.

Then, we inquired whether the lack of the internal NPT2A ^494^TRL^496^ PDZ motif affects hormone-sensitive phosphate transport. FGF23- or PTH-inhibitable phosphate uptake was measured in CRISPR/Cas9 Npt2a knockout OK cells transfected with NPT2A with the triple Ala replacement at the internal ^494^AAA^496^or CT-AAA PDZ motif and compared to WT NPT2A. Unexpectedly, ablating the internal PDZ motif prevented FGF23- or PTH-sensitive phosphate uptake compared to WT NPT2A but not basal phosphate transport (**Figure 3**).

**Figure 3.**
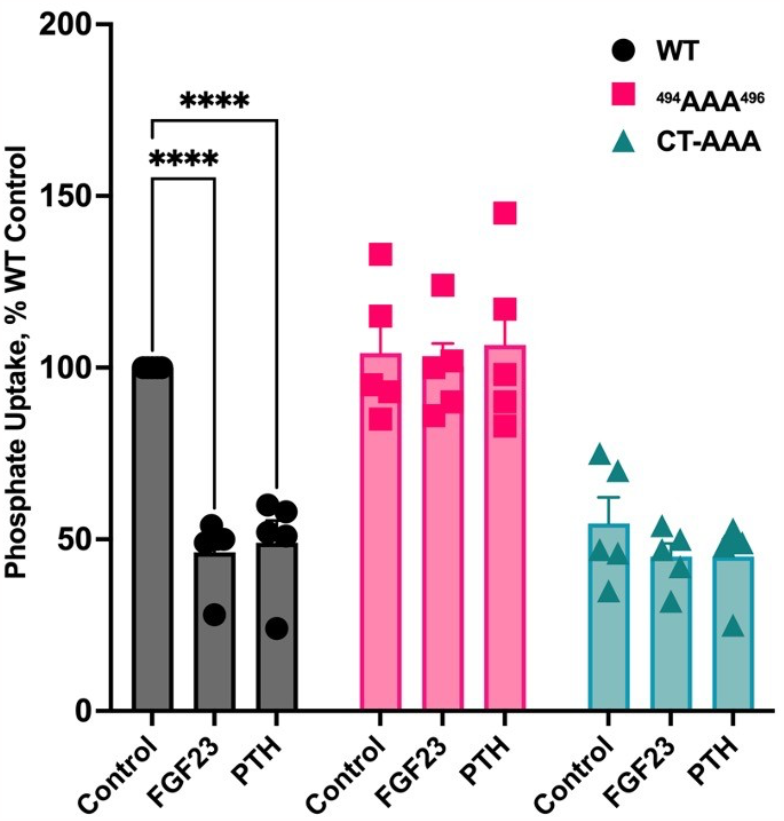
Triple alanine replacement in NPT2A internal or carboxy-terminal PDZmotif inhibits hormonesensitive phosphate uptake. NPT2A ^494^AAA^496^ and CT-AAA PDZ ligand mutants do not support FGF23- or PTH-sensitive phosphate uptake. CRISPR/Cas9 Npt2a knockout OKcells were transfected with the indicated NPT2A mutants. Phosphate uptake was assayed in the presence or absence of FGF23 or PTH, as indicated. *n*=5. ****p<0.0001PTH or FGF23 *vs* control.

In contrast, the NPT2A CT-AAA variant reduced resting phosphate transport (Figure 3). Consistent with the phosphate uptake experiments, the NPT2A ^494^AAA^496^ mutant was immunoprecipitated with NHERF1, whereas NPT2A CT-AAA did not (**Figure 4A**). **Figure 4B** displays the quantification of NPT2A ^494^AAA^496,^ immunoprecipitation with NHERF1 comparable to WT NPT2A. The spatial localization of NPT2A variants was analyzed by confocal fluorescence microscopy.

**Figure 4.**
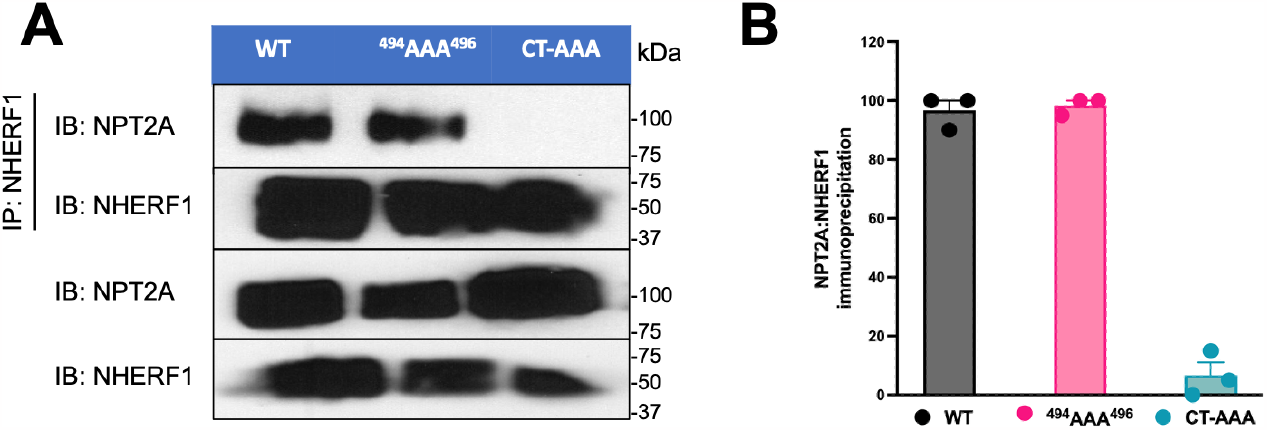
Effect of triple alanine replacement in NPT2A on the binding with NHERF1. (A) A representative experiment depictingNPT2A immunoprecipitating with NHERF1. The NPT2A ^494^AAA^496^ mutant immunoprecipitated with NHERF1, whereas NPT2A CT-AAA does not. Immunoblots for IP are shown in the top two panels. Lysates controls are in the bottom panels. Molecular weight markers (kDa) are to the right of the blots. (B)Quantification of NPT2A immunoprecipitated with NHERF1. CRISPR/Cas9Npt2a knockout OK cells were transfected with HA-GFP-NPT2A WT, ^494^AAA^496^or CT-AAA and FLAG-NHERF1. The amount of mutant NPT2Aimmunoprecipitating with NHERF1 was normalized to the amount of WT-NPT2A, which was defined as 100%. *n*=3, **** p<0.0001 CT-AAA *vs*. WT.

WT NPT2A, NPT2A ^494^AAA^496,^and NHERF1 are expressed at the apical membrane under resting conditions in the absence of PTH (**Figure 5A, B**). WT NPT2A and NPT2A ^494^AAA^496^colocalize broadly with NHERF1 but not with NPT2A CT-AAA (**Figure 5C**). Upon PTH treatment, WT NPT2A internalized extensively in the presence of NHERF1 (Figure 5A), whereas NPT2A ^494^AAA^496^remained localized at the apical membrane (Figure 5B). NPT2A CT-AAA was essentially unchanged (Figure 5C). Colocalization of NPT2A variants with NHERF1 was quantified as described in **Materials and Methods** and presented in **Figure 5D**.

**Figure 5.**
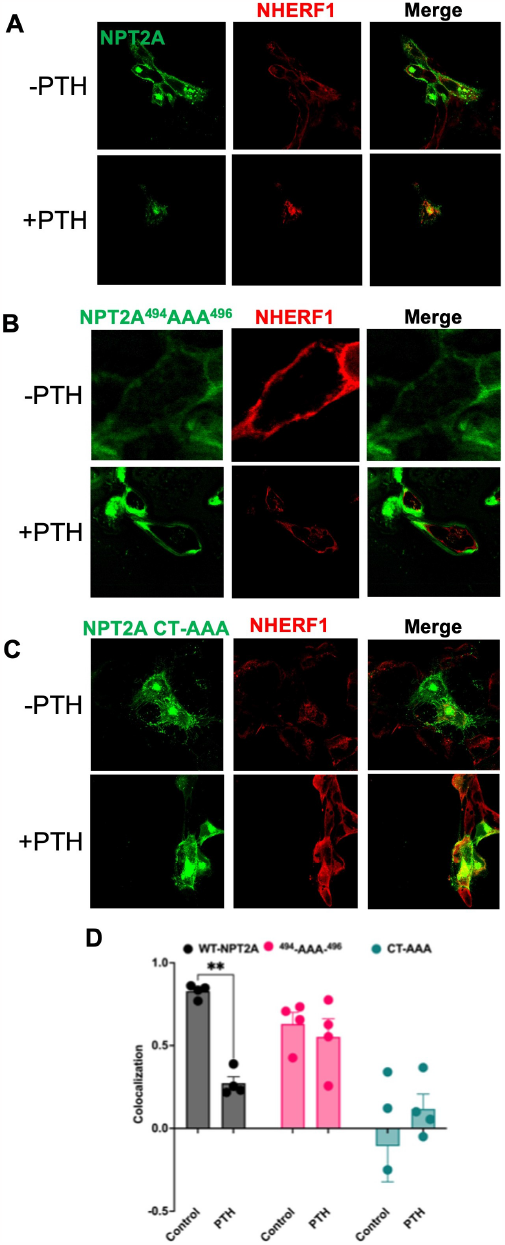
Confocal fluorescence localization of NPT2A (green) with NHERF1 (red) inCRISPR/Cas9 Npt2a knock-out OK cells. Under resting conditions in the absence of PTH (upper row), (A)WT NPT2A or (B) NPT2A ^494^AAA^496^ and NHERF1 areexpressed at theapicalmembrane. Merged imagesareshown on therightand indicateextensive Colocalization for WT NPT2A or NPT2A ^494^AAA^496^with NHERF1 but not for (C) NPT2A CT-AAA. Upon addition of PTH (bottom row), (A) WT NPT2A or (B) NPT2A ^494^AAA^496^internalized extensively in the presence of NHERF1, but (C) NPT2A CT-AAA was essentially unchanged. (D) Colocalization was quantified using Image J. The calculated threshold overlap score (TOS) between +1 and -1 indicates Colocalization between NPT2A variants and NHERF1 or its absence, respectively. *n*=4.

Together, our experimental data demonstrate that the internal NPT2A^494^TRL^496^ PDZ motif is required for FGF23- and PTH-triggered dissociation of NPT2A from NHERF1, while the NPT2A CT-TRL PDZ ligand determines membrane localization of NPT2A.

### Disease-associated mutations in the NPT2A internal PDZ motif abolish hormone-sensitive phosphate transport

Mutation of the highly conserved Arg^495^ in the internal NPT2A ^494^TRL^496^ motif to Cys or His is associated with congenital elevated renal phosphate excretion and hypophosphatemia in children [21, 22]. These observations suggest a more precise role for the cryptic internal PDZ motif than revealed by the experimental triple AAA construct described above. We, therefore, introduced NPT2A variants, where Arg^495^was replaced by Cys or His, i.e., the reported human mutations. Phosphate uptake was measured in CRISPR/Cas9 Npt2a knockout OK cells transfected with the NPT2A Arg^495^Cysor Arg^495^Hisvariant. Notably, FGF23- and PTH-sensitive phosphate uptake was abolished by the disease-associated NPT2A variants (**Figure 6**).

**Figure 6.**
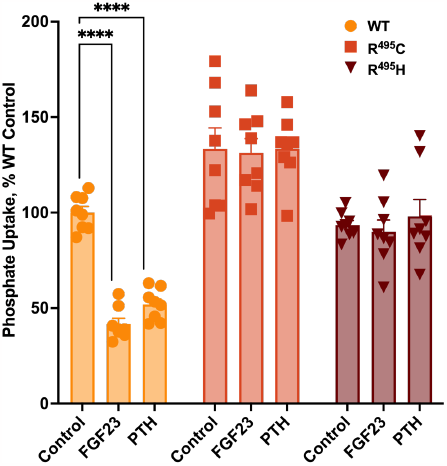
Disease-associated mutations in the NPT2A internal PDZ ligand block FGF23- or PTH-sensitive phosphate transport. Arg^495^(R^495^) in the ^494^TRL^496^ motif was substitutedby Cys (R^495^C) or His (R^495^H). CRISPR/Cas9 Npt2aknockout OK cells were transfected with the indicated NPT2A variants. Phosphate uptake was assayed in the presence or absence of FGF23 or PTH, as indicated. *n*=8,**** p<0.0001 PTH or FGF23 vs. control.

Coimmunoprecipitation experiments revealed that both NPT2A Arg^495^Cysand Arg^495^Hisvariants associate with NHERF1 under resting conditions. However, compared to WT NPT2A, neither Arg^495^Cysnor Arg^495^Hisdissociates from NHERF1 upon challenge with PTH (**Figure 7A**), the standard biological action that causes cessation of phosphate transport. Concurrently, as demonstrated by confocal fluorescence microscopy, NPT2A Arg^495^Cys and Arg^495^His variants colocalized with NHERF1 at apical cell membranes like WT NPT2A (**Figure 7B, C, D** [left panel]). Consistent with the functional results, NPT2A Arg^495^Cys and Arg^495^His variants do not internalize in response to PTH but remain at the apical membrane (**Figure 7C, D** [right panel]). Quantified NPT2A:NHERF1 complex Colocalization illustrates that, in contrast to WT NPT2A, the disease variants do not dissociate upon challenge with PTH (**Figure7E**).

**Figure 7.**
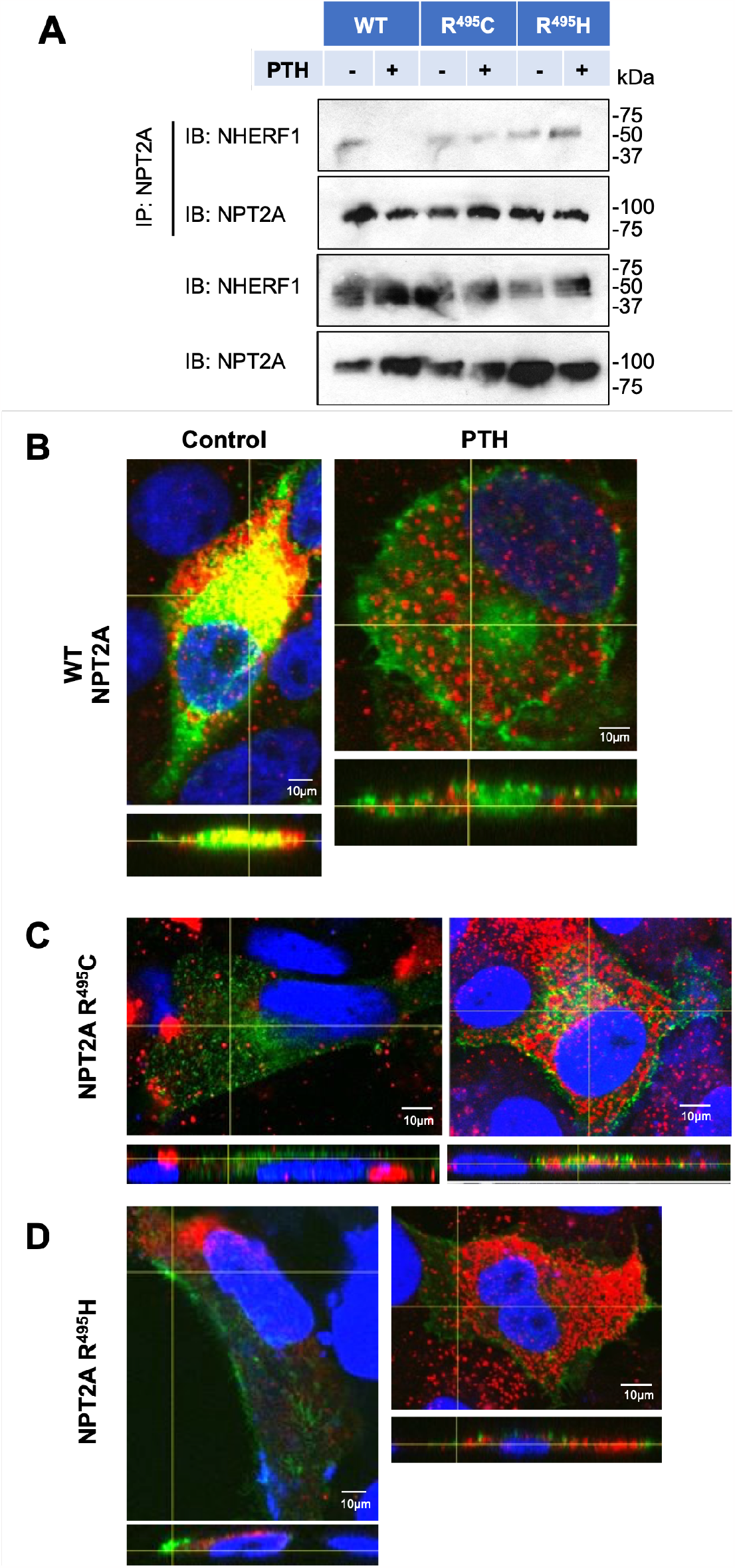
Disease-associated mutations in the NPT2A internal PDZ ligand do not interfere with NPT2A-NHERF1 association or NPT2A-NHERF1 colocalization. (A) HA-GFP-NPT2A R^495^Cor R^495^H do not dissociate from NHERF1 in response to PTH in OKcells compared to WT NPT2A. OK cells were transfected with HA-GFP-NPT2A (WT, R^495^C, or R^495^H) and FLAG-NHERF1. (B) Confocal fluorescence colocalization of WT NPT2A, (C) NPT2A R^495^C, or (D) NPT2AR^495^H (green) with NHERF1 (red) in OK cells. (E) NPT2A:NHERF1 colocalization was quantified. We calculated the Threshold Overlap Score (TOS)as a measure for Colocalization for each NPT2A mutant using confocal microscopy images and Image J. *n*=4, ****p<0.0001 PTH *vs*. control.

Class I PDZ domains, such as those in NHERF1, favor ligands containing Thr or Ser at position -2 relative to the carboxy-terminus, position 0 [24]. Phospho-resistant modifications of Ser/Thr at residue -2 of class I PDZ-recognition motifs impair the binding [25, 26]. Thr^494^,located at position -2 of the internal NPT2A ^494^TRL^494^ PDZ motif (T^*−*2^R^*−*1^L^0^), serves as a specific determinant of a canonical PDZ-ligand interaction. We hypothesized that replacing Arg^495^ by Cys or His of the internal NPT2A ^494^TRL^496^ motif impairs the phosphorylation of Thr^494^. To test this idea, HA-GFP-NPT2A Arg^495^Cys or Arg^495^His variants were treated with PTH in transfected cells and immunoblotted with a custom antiphospho-^494^TRL^496^antibody **(Figure 8A**) and for total NPT2A (**Figure 8B**). Representative immunoblots (**Figure 8A, B**) and quantification (**Figure 8C**) clearly show that PTH promotes phosphorylation of Thr^494^ in WT NPT2A but not in NPT2A Arg^495^Cys or Arg^495^His variants.

**Figure 8.**
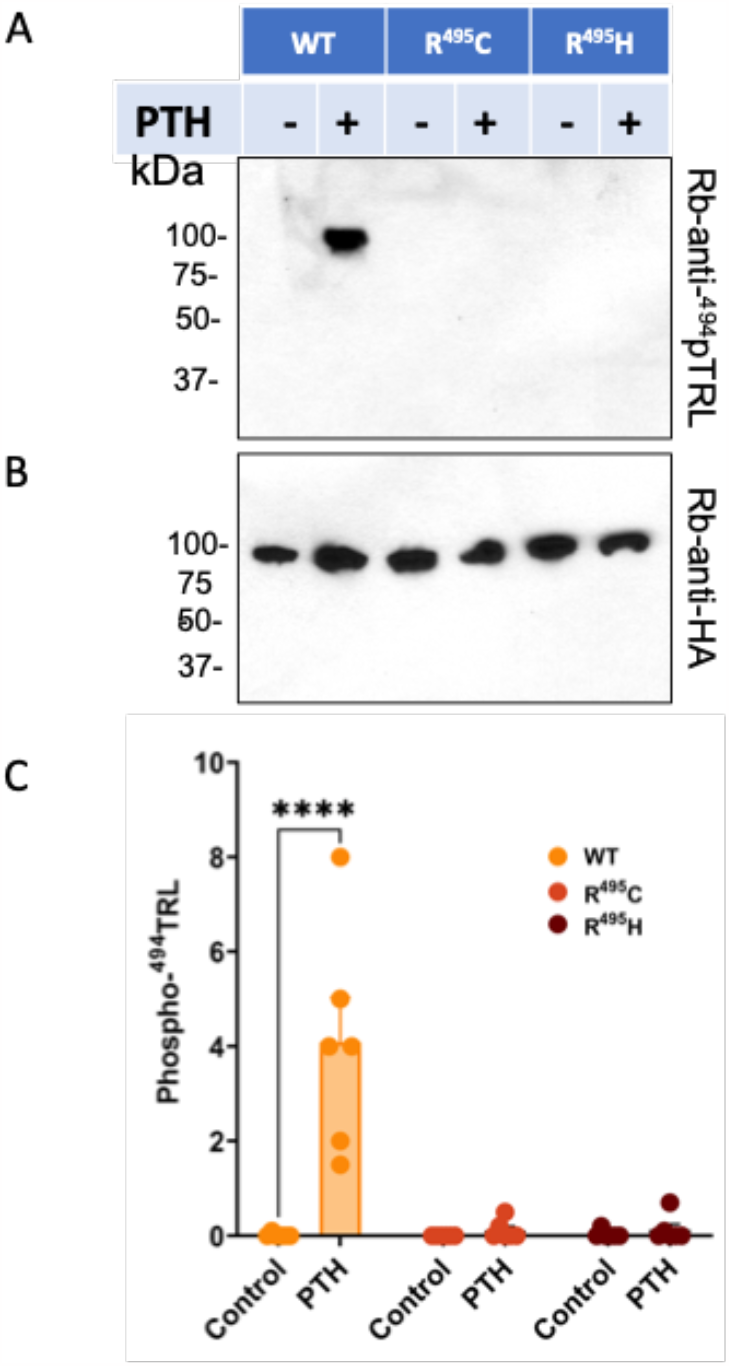
Replacement of Arg^495^ by Cys or His in NPT2A internal TRLmotif prevents phosphorylation of the upstream Thr^494^ compared to WT. Representative immunoblots of HA-GFP-NPT2A Arg^495^Cys or Arg^495^His variants treated with PTH and immunoblotted for (A) anti-p^494^TRL^496*−*^antibody or (B) total NPT2A. Molecular weightmarkers (kDa) are shown on the left of the blots. (C) Quantification of immunoblot analysis. *n*=3. ****p<0.0001 PTH *vs*. control.

Next, we inquired whether Thr^494^ is involved in hormone-sensitive phosphate transport. Phosphate uptake was measured in CRISPR/Cas9 Npt2a knockout OK cells transfected with an NPT2A construct, where Thr^494^was replaced conservatively by Cys (Thr^494^Cys), and the cells were treated with FGF23 or PTH. Notably, the NPT2A Thr^494^Cys mutant impaired the inhibitory response to FGF23 and PTH (**Figure 9A**) but did not affect resting phosphate uptake. We then turned to the interaction of the NPT2A Thr^494^Cys construct with NHERF1.

**Figure 9.**
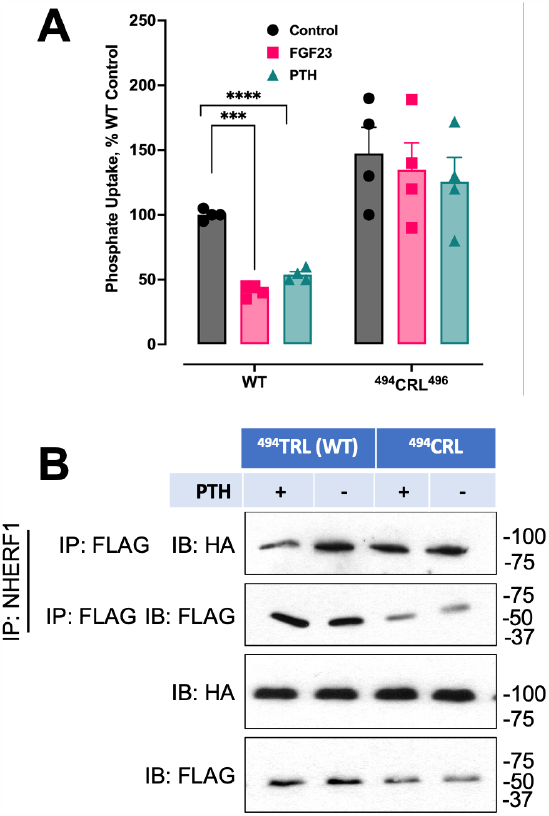
Thr^494^/Cys replacement in the internal NPT2A PDZ ligand blocks hormone-sensitive phosphate uptake. (A) NPT2A ^494^CRL mutant is refractory to PTH and FGF23 action on phosphate uptake. CRISPR/Cas9Npt2aknockout OK cells were transfected with the indicated NPT2A mutant and treated with FGF23 or PTH. n=4. *** p<0.001FGF23 vs control; ****p<0.0001 PTH vs control; * p<0.1 FGF23 *vs* PTH. (B) Thr^494^/Cyssubstitution supports the binding of HA-NPT2A and FLAG-NHERF1 but prevents the dissociation of HA-NPT2A from FLAG-NHERF1 in the presence of PTH compared to WT HA-NPT2A as assessed by representative immunoblot. Molecular weight markers (kDa) on the right of blots. *n*=4.

Representative immunoblots presented in **Figure 9B** demonstrate that HA-NPT2A Thr^494^Cys binds FLAG-NHERF1. Notably, the replacement of Thr by Cys prevented the dissociation of HA-NPT2A from FLAG-NHERF1 upon treatment with PTH (**Figure 9B**). Then CRISPR/Cas9 Npt2a knockout OK cells were transfected with HA-GFP-NPT2A Thr^494^Cys or WT NPT2A, treated with PTH, and immunoblotted for Rb-anti-^494^pTRL^496^antibody, or Rb-anti-HA. As expected, NPT2A Thr^494^Cys does not phosphorylate in response to PTH (**Figure 10**).

**Figure 10.**
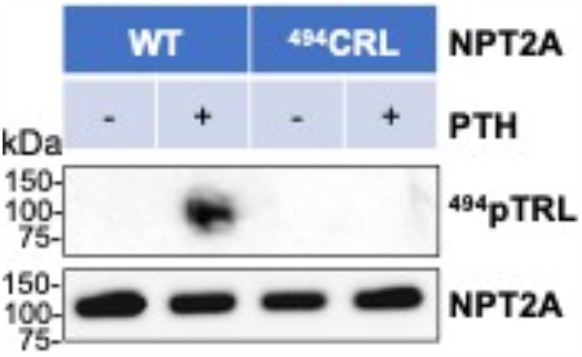
PTH does not promote the phosphorylation of NPT2A Thr^494^Cys. Representative immunoblot of HA-GFP-NPT2A (WT) or NPT2A Thr^494^Cys (CRL) in CRISPR/Cas9 Npt2a knockout OK cells treated with PTH and immunoblotted with Rb-anti-^494^pTRL^496^antibody (top panel), or total NPT2A (bottom panel). Molecular weigh tmarkers (kDa) are shown on the left of the blots. *n*=3.

It is enigmatic how Arg^495^ mutation affects phosphorylation of the vicinal Thr^494^. We speculated that the binding of NPT2A and NHERF1 PDZ2 is necessary for the phosphorylation of Thr^494^ and that the Arg^495^mutation interferes with Thr^494^phosphorylation. OKH cells were transfected with WT-HA-GFP-NPT2A and FLAG-NHERF1 or FLAG-NHERF1 S2 (PDZ1), treated with PTH and immunoblotted for Rb-anti-^494^pTRL^496^antibody, Rb-anti-HA, or Rb-anti-FLAG (**Figure 11)**. Phosphorylation of Thr^494^ was detected in cells transfected with WT NHERF1 but not NHERF1 S2, illustrating the requirement for NHERF1 PDZ2 **(Figure 11**).

**Figure 11.**
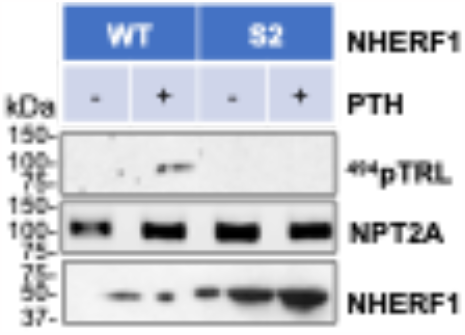
PTH-induced phosphorylation of NPT2A Thr^494^ requires NHERF1. Representative immunoblots of HA-GFP-NPT2A and FLAG-WT NHERF orFLAG-NHERF1 S2 in OKH cells treated with PTH and immunoblotted for Rb-anti-p^494^TRL^496^antibody (top panel), Rb-anti-HA (middle panel), or Rb-anti-FLAG (bottom panel). Molecular weight markers (kDa) are shown on the left of the blots. n=*3*.

### Molecular modeling of the internal NPT2A^494^TRL^496^ PDZ motif

The structure of NPT2A has not been solved. Therefore, we applied AlphaFold2 (AF2) [27] to generate a 3D computational structure of NPT2A. To explore residue-residue interactions specific for the NPT2A internal ^494^TRL^496^ PDZ motif coordinates of residues 310-536 were used to establish a model for MD simulation. These coordinates correspond to the transmembrane helices 3 to 7 (TM3-TM7) as assigned elsewhere [28]. All-atom explicit solvent MD simulations (200-ns) were performed using AMBER16 [29]. Simulation details are provided in the Materials and Methods. The structure resulting from the MD simulation is similar to that predicted by AF2. The RMSD values calculated for the backbone atoms relative to the AF2 structure along the equilibration and production simulations remain stable for *α*-helices (not presented here). In contrast, a loop connecting TM5b and TM6a (residues 453-482) changes its conformation along the MD simulation. The N-terminal flanking region (residues 507-536, segment of TM7) is mobile. These regions AF2 may predict with low confidence compared to *α*-helices [30]. A representative snapshot from the MD simulation demonstrates that the sidechains of Thr^494^ and Leu^496^ are oriented to the solvent. In contrast, the sidechain of Arg^495^ turns opposite toward the core of NPT2A (**Figure 12**). Calculation of hydrogen bonds (cpptraj AMBER16{Case, 2016 #25}) [29] along the production simulation suggests that Arg^495^ (TM6) forms a salt bridge with Glu^438^ (TM5a) (**Figure 12**).

**Figure 12.**
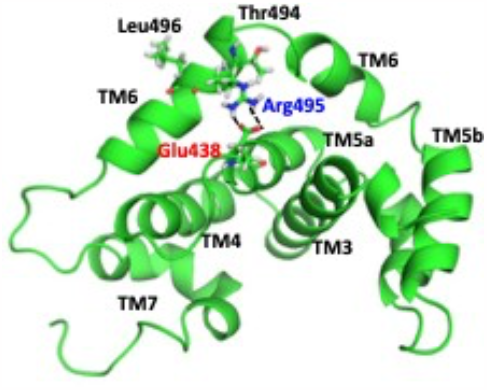
A computational model of NPT2A (395-536 aa) from MD simulation. A representative snapshot showing the orientation of the NPT2A internal ^494^TRL^494^PDZ motif. Thr^494^ and Leu^496^ are solvent-exposed, whereasArg^495^ is buried within the pocket created by TM5a, TM6a, and TM6b. A predicted salt bridge between Arg^495^ (TM6b) and Glu^438^(TM5a) is presented as a dotted black line. The TM helices in the MD model were assigned as published elsewhere[28].

We computationally replaced Arg^495^ with Cys or His and conducted additional MD simulations (100-ns). The equilibration and production simulations were performed as described for WT NPT2A. MD simulations predict that the replacement of Arg^495^ by Cys or His does not induce conformational changes but rather abrogates the electrostatic interaction with Glu^438^. The side chain of Thr^494^ or Leu^496^ does not change its orientation and remains solvent-exposed.

The computational study supports our experimental finding, thus indicating that AF2 can reliably assess the accuracy of NPT2A structure from amino acid sequences and, in combination with MD simulation, is extremely useful.

## DISCUSSION

The Na^+^-dependent phosphate cotransporter-2A (NPT2A, SLC34A1) is the principal regulator of phosphate homeostasis. Its most notable structural element is its carboxy-terminal PDZ ligand that binds Na^+^/H^+^ Exchanger Regulatory Factor-1 (NHERF1, SLC9A3R1), a multidomain PDZ protein that determines apical NPT2A localization, is required for phosphate transport. NPT2A also harbors an undescribed internal identical PDZ motif ^494^TRL^496^. The objectives of the present work were to determine what, if any, biological function the internal PDZ motif fulfills. An important clue suggesting a potentially important role emerged from two case reports describing hypophosphatemia in children with inherited mutations located within the internal PDZ motif, though it was not initially appreciated. A third goal was to resolve the biochemical riddle of how Arg mutation prevents obligate, hormone-triggered phosphorylation of the adjacent Thr.

To characterize the two PDZ motifs, we generated NPT2A mutants where CT-TRL or ^494^TRL^496^ was replaced with AAA. We show here, using immunoblot and immunoprecipitation analysis and confocal fluorescence imaging, that CT-AAA disrupts the interaction with NHERF1 (**Figure 4**) and, concurrently, decreases the distribution of NPT2A at the apical membrane and renders NPT2A refractory to PTH (**Figure 5C**). We conclude that the NPT2A CT-TRL defines the apical membrane colocalization of NPT2A with NHERF1 that is an antecedent and required component for hormone-sensitive phosphate transport.

Coimmunoprecipitation experiments indicated that the internal NPT2A ^494^TRL^496^ PDZ motif binds NHERF1 PDZ2 (**Figure 2**), which we consider a regulatory domain. In contrast to CT-TRL, ablation of the internal NPT2A ^494^TRL^496^ motif does not change NPT2A apical localization (**Figure 5B**) or the interaction with NHERF1 (**Figure 4**). Intriguingly, NPT2A ^494^AAA^496^ blocks the inhibition of phosphate uptake by FGF23 or PTH but does not affect resting conditions (**Figure 3**). The ^494^TRL^496^ motif is, therefore, an essential regulator of hormone-triggered phosphate transport.

Two recent clinical reports describe hereditary hypophosphatemia in children harboring Arg^495^His or Arg^495^Cys variants within the internal NPT2A PDZ motif [21, 22]. It was not appreciated that these disease-causing mutations were in a canonical, if cryptic, PDZ motif. We show here that the two variants, NPT2A Arg^495^Cys and Arg^495^His, block inhibition of phosphate uptake in response to FGF23 or PTH (**Figure 6**) while fully able to complex with NHERF1 (**Figure 7A**). In agreement with immunoblot analysis, NPT2A Arg^495^His or Arg^495^Cys fail to internalize in response to PTH and remain at apical cell membranes (**Figure 7C, D**). The data raise the question of why mutating Arg^495^ to Cys or His [21, 22] at position -1 of the PDZ motif, which is considered permissive [24], blocks phosphate transport (**Figure 6**). In the absence of NPT2A structural information, we used an AF2 predictor to generate a computational model to resolve this enigma. Further MD simulations show that Thr^494^ and Leu^496^ are solvent-exposed and may be involved in the interactions with NHERF1. In contrast, Arg^495^forms a salt bridge with Glu^438^, thereby connecting TM6 and TM5a (**Figure 12**). We speculate that Thr^494^ (position -2) and Leu^496^(position 0) are PDZ ligand-binding determinants involved in the interaction with NHERF1 PDZ2, whereas Arg^495^ (permissive position -1) is essential for post-translational modification. The MD simulation predicts that the substitution of Arg^495^by Cys or His disrupts the salt bridge between Arg^495^ and Glu^438^ and changes the electrostatics that may prevent phosphorylation of the upstream Thr^494^. We experimentally generated an NPT2A construct to probe this idea, where Cys replaced Thr^494^. We assumed that the similarity between the side chain of Cys (SH) and Thr (OH) would permit Class I PDZ-ligand interaction [24] and not impede the association with NHERF1. At the same time, Cys cannot be phosphorylated compared to Thr and would prevent PTH action. This was the case. Indeed, the NPT2A Thr^494^Cys variant binds NHERF1 (Figure 9B), is not phosphorylated (Figure 10), and does not support PTH-sensitive inhibition of phosphate uptake (Figure 9A). By employing a custom anti-^494^pTRL^496^ antibody, we confirmed the phosphorylation of Thr^494^ after PTH treatment (Figure 11). Of note, phosphorylation of NPT2A Thr^494^ was detected only in cells transfected with NHERF1. Furthermore, phosphorylation of Thr^494^ was not detected in the absence of NHERF1 PDZ2 (S2 construct) (Figure 11), supporting our contention that the NPT2A internal PDZ ligand binds to NHERF1 PDZ2. We speculate that the association between the internal NPT2A^494^TRL^496^ PDZ motif and NHERF1 PDZ2 is necessary for PTH-induced phosphorylation of the Thr^494^. Additional studies will further characterize this phenomenon.

In summary, we identify and characterize the NPT2A internal^494^TRL^496^ PDZ motif as a novel, heretofore undescribed determinant of hormone-sensitive phosphate transport. We propose an advanced model where both NPT2A PDZ binding motifs play distinct roles in basal and hormone-sensitive phosphate regulation: the carboxy-terminal PDZ ligand defines the apical localization of NPT2A through the interaction with NHERF1. The internal PDZ motif is an essential determinant of hormone-triggered action on phosphate transport and NPT2A trafficking.

## Materials and Methods

### Peptides

Human [Nle^8,18^,Tyr^34^]PTH(1–34) was purchased from Bachem (H9110). Recombinant human R179Q-FGF23(25–251) (referred to henceforth as FGF23), which is resistant to furin cleavage and inactivation, was obtained from R&D Systems (2604-FG-025).

### Antibodies

Antibodies Rabbit polyclonal anti-NHERF1 (ab3452) and monoclonal anti-NHERF1 (Abcam ab31111) were purchased from Abcam. M2-mouse monoclonal anti-HA-agarose (A2095) and rabbit polyclonal anti-FLAG antibodies (F7425) were purchased from Sigma. Rabbit polyclonal anti-HA (sc-805) was purchased from Santa Cruz. Horseradish Peroxidase-conjugated Goat Anti-Rabbit (Dako P044801-2) and Rabbit Anti-Mouse (Dako P044701-5) secondary antibodies were purchased from Agilent. Primary antibodies were used at a 1:1000 dilution for immunoblots; secondary antibodies were employed at a 1:5000 dilution. The agarose-conjugated anti-HA was used at a 1:50 dilution for immuno-precipitation experiments.

### Cell culture

A new CRISPR/Cas9 Npt2a knockout OK cell line was developed to explore human NPT2A structural determinants involved in binding with NHERF1 and hormone-sensitive phosphate transport. FGF23- and PTH-sensitive phosphate uptake in these cells was rescued with human HA-GFP-NPT2A or mutant variants (^494^AAA^496^, CT-AAA, Arg^495^Cys, Arg^495^His, and ^494^CRL^496^).

Opossum kidney (parental OK or NHERF1-deficient OKH) cells [9] were grown in DMEM/Ham’s F-12 50:50 medium (Mediatech, 10-090-CV) supplemented with 10% FBS and 1% pen/strep. Cells were transfected with the indicated plasmids using FuGENE 6 (Promega) or Lipofectamine 3000 (Invitrogen) according to the manufacturer’s instructions. Stable cells expressing FLAG-NHERF1 or FLAG-NHERF1 constructs (S1, S2, or S1S2) were prepared by screening with puromycin or G418.

### Preparation of endogenous NHERF1 and NHERF1 constructs

WT or NHERF1 S1, S2, or S1S2 constructs tagged with FLAG epitope were transfected using Lipofectamine 3000 (Invitrogen) as described previously [11].

### Immunoprecipitation

OK cells were transfected with HA-GFP-WT-NPT2A (WT) or HA-GFP-NPT2A-^494^CRL^496^ (^494^CRL^496^) and FLAG-NHERF1. After 48 h post-transfection, cellswereserum-starved overnightand then treated for 2 h in thepresence of 100 nM PTH. Lysates were prepared, and NHERF1 was immunoprecipitated with Ms-anti-FLAG agarose (Sigma). NPT2A was immunoblotted with Rb-anti-FLAG (Sigma). NHERF1 was immunoblotted using Rb-anti-HA (Santa Cruz).

Acustomanti-phosphoNPT2AThr^494^ (^494^pThr)antibody(10343GenScript)generatedagainsttheNPT2A ^485^ILLWYPVPCT^494^R sequence was applied to verify phosphorylation of Thr^494^ as before [12]. CRISPR/Cas9 Npt2a knockout OK cells were transfected with HA-GFP-NPT2A (WT) or NPT2A with the triple AAA replacement (^494^AAA^496^) for 24 h, and serum-starved overnight. The next day, cells were treated with 100 nM PTH for 2 min. Lysates were prepared in the presence of phosphatase inhibitors. Cells were immunoblotted for ^494^pTRL or total NPT2A (Novus). Primary antibody (1:1000) or Goat anti-rabbit-HRP secondary antibody (1:5000) were applied for 10 min.

### Phosphate uptake

CRISPR/Cas9 Npt2a knockout OK cells transfected with HA-GFP-NPT2A or NPT2A mutant variants were seeded on 12-well plates. When the cells reached confluence (2-3 days after passaging), they were treated with 100 nM PTH(1&#8209;34) or FGF23 in cell culture media. After 2 h, the hormone-supplemented media was aspirated, and the wells were washed 3 times with 1 mL of Na-replete wash buffer (140 mM NaCl, 4.8 mM KCl, 1.2 mM MgSO_4_, 0.1 mM KH_2_PO_4_, 10 mM HEPES, pH 7.4). The cells were incubated with 1 *μ*Ci of ^32^P orthophosphate (PerkinElmer, NEX053) in 1 mL of Na-replete wash buffer for 10 min. Phosphate uptake was terminated by placing the plate on ice and rinsing the cells 3 times in Na-free wash buffer (140 mM N-methyl-D-glucamine (NMDG), 4.8 mM KCl, 1.2 mM MgSO_4_, 0.1 mM KH_2_PO_4_, 10 mM HEPES, pH 7.4). The cells in each well were extracted using 500 *μ*l of 1% Triton X100 (Sigma Cat X100) in water overnight at 4°C. A 250-*μ*l aliquot was processed for analysis on a Beckmann Coulter LS6500 liquid scintillation counter. Data were normalized, where 100% was defined as the CPM of phosphate uptake under control conditions.

### Confocal microscopy

Confocal fluorescence imaging was performed as described previously with some modifications [9]. Briefly, CRISPR/Cas9 Npt2a knockout OK cells or OK cells, as indicated, were seeded on collagen-coated coverslips. 24 h later, the cells were transiently transfected with HA-GFP-NPT2A (WT or mutant constructs, as shown in Figures 5A, B, C, and 7B, C, D), and FLAG-NHERF1. 48 post-transfected, the cells were serum-starved overnight and treated with Vehicle (control) or 100 nM PTH for 2 h, fixed in 4% paraformaldehyde, and stained with Rb-anti-FLAG (Sigma) primary antibody (1:50) and Goat anti-rabbit-Alexa546 (Invitrogen) secondary (1:1000) antibody. Coverslips were mounted for immunofluorescence microscopy using ProLong™ Gold Antifade Mountant with DAPI (Thermofisher P36941) and analyzed using an Olympus FluoView 1000 microscope with a 63x oil immersion objective. Colocalization analyses for HA-GFP-NPT2A variants and FLAG-NHERF1 were performed with ImageJ. Confocal images were quantified using the EzColocalization plugin for ImageJ and calculating the Threshold Overlap Score (TOS) [31]. TOS ranges from +1 (perfect correlation) to -1 (perfect but negative correlation), with 0 denoting the absence of a relationship. Confocal fluorescence imaging was also performed for NPT2A with the disease-associated Arg^495^ Cys and Arg^495^ His NPT2A variants as described above.

### System setup and molecular dynamics simulation

AlphaFold2 [27] was applied to create a 3D model of full-length NPT2A. Because of computationallimitations, the model for MD simulation includes TM4-TM7 (residues 310-536). Files for simulation were prepared using the Leap module of AMBER16 [29]. The WT NPT2A model was solvated with TIP3P water molecules [33] in a periodically replicated box and neutralized with Cl^*−*^ ions. The structure was relaxed by 100 cycles of steepest descent followed by 400 cycles of conjugate gradient energy minimization procedure using the AMBER16 pmemd module [29] with harmonic restraints (force constant - 0.5 kcal/mol/Å2) applied to all protein atoms. A 1-ns MD simulation run was performed under the NPT ensemble [constant number of particles (N), pressure (P), and temperature (T)] to reach the correct density of liquid water (∼ 1 g/mL). During this equilibration, harmonic restraints were applied to all residues and methodically lowered from k_*s*_ = 10 kcal/mol/Å^2^to 0.1 kcal/mol/Å^2^. The simulation was continued under the NVT ensemble [constant number of particles (N), volume (V), and temperature (T)] at 300 K with harmonic restrains k_*s*_ = 0.1 kcal/mol/Å^2^ applied to the N-terminal and C-terminal backbone atoms to prevent diffusion of the protein from the simulation box. Langevin coupling algorithm was applied. The energy minimization and MD simulation were carried out using the AMBER16 package with the AMBER force field (ff99SB). Since the starting structure was not experimentally resolved conformation, we performed a long equilibration protocol (50 ns). A production simulation was run for approximately 150 ns with the temperature at 300 K and constant volume with the same harmonic restraints as in the equilibration phase. Coordinates were saved every 2 ps for later analyses. The NPT2A Arg^495^Cys or Arg^495^His model was built by replacing Arg^495^with Cys or His, respectively, using the Leap module of AMBER16 [29]. Except for the length, the simulation details, equilibration, and production simulations were set up as described for WT NPT2A. After solvation and ionization, each system has a total of ∼32920 atoms. The equilibration and production simulations of wild-type and mutant systems were monitored by calculating the root-mean-square deviation (RMSD) (cpptraj of AMBER16) [29] of the backbone atoms from their initial positions (not presented here). The hydrogen bond analysis was performed using the ccptraj module of AMBER16 [29].

### Statistical analysis

Results were analyzed using Prism 9 software. Data represent the mean *±* SD or SEM calculated on at least three independent experiments. Each figure’s legend indicates the exact number of experiments performed and used for statistical analysis. p Values were calculated using one-way ANOVA with the Brown-Forsythe test, two-way ANOVA with Šidák’s or Dunnett’s multiple comparison test, or three-way ANOVA with Turkey’s multiple comparison procedure. p Values were depicted by *p<0.05, **p<0.01, ***p < 0.001, and ****p<0.0001. p Values <0.05 were considered statistically significant.

## Data availability

The authors agree to make any materials, data, and associated protocols available upon request.

## Competing Interests

The authors declare no conflicts of interest in connection with this article.

### CRediT Contributions

Peter A. Friedman: Conceptualization, Formal analysis, Writing — original draft, Writing — review and editing, Visualization, Supervision, Project administration, Funding acquisition

Tatyana Mamonova: Conceptualization, Formal analysis, Investigation,

Writing — original draft, Writing — review and editing.

W. Bruce Sneddon: Investigation, Methodology, Resources, Writing - Original Draft, Writing - Review & Editing, Visualization

## Abbreviations

NPT2A: Na^+^-dependent phosphate cotransporter
NHERF1: Na^+^/H^+^ Exchanger Regulatory Factor-1
PTH: parathysroid hormone
FGF23: Fibroblast growth factor 23
OK(H): opossum kidney cells
AF2: AlphaFold2
TM: transmembrane domain
MD simulation: molecular dynamics simulation
RMSD: root-mean-square deviation
TOS: Threshold Overlap Score

## Acknowledgments

We thank Dr. Qiangmin Zhang for preparing the NPT2A constructs used here.

